# *Arabidopsis ABIG1* Functions in Laminar Growth and Polarity Formation through Regulation by *REVOLUTA* and *KANADI*

**DOI:** 10.1101/2021.06.16.448748

**Authors:** Jesus Preciado, Kevin Begcy, Tie Liu

## Abstract

Leaf laminar growth and adaxial-abaxial boundary formation are fundamental outcomes of plant development. Boundary and laminar growth coordinate the further patterning and growth of the leaf, directing the differentiation of cell types within the top and bottom domains and promoting initiation of lateral organs along their adaxial/abaxial axis. Leaf adaxial-abaxial polarity specification and laminar out-growth are regulated by two transcription factors, *REVOLUTA* (*REV*) and *KANADI* (*KAN*). *ABA INSENSITIVE TO GROWTH 1* (*ABIG1*) is a *HOMEODOMAIN-LEUCINE ZIPPER* (*HD-ZIP*) Class II transcription factor and is a direct target of the adaxial-abaxial regulators *REV* and *KAN*. To investigate the role of *ABIG1* in the leaf development and establishment of polarity, we examined the phenotypes of both gain-of-function and loss-of-function mutants. Through genetic interaction analysis with *REV* and *KAN* mutants, we have determined that *ABIG1* plays a role in leaf laminar-growth as well as in adaxial-abaxial polarity establishment. Genetic and physical interaction assays showed that *ABIG1* interacts with the transcriptional corepressor *TOPLESS* (*TPL*). This study provides new evidence that another *HD-ZIP II* gene, *ABIG1*, facilitates growth through the corepressor *TPL*.

**Highlight:** ABIG1, a HD-ZIP Class II transcription factor, promotes laminar growth and adaxial-abaxial polarity through the regulation of *REV* and *KAN*.

## Introduction

Plant architecture is one of the basic tenets of plant biology. The initiation of the spatial arrangement of root, shoot, and floral meristems is an elaborate, intricate and well-coordinated process. The ‘body plan’ of shoot architecture begins with the establishment of a primary axis of growth (Leyser, 2009). Abaxial and adaxial characteristics can be defined in the shoot apical meristem (SAM), the leaves, and the vasculature (Byrne, 2006). In the SAM, the central zone is the adaxial region and contains pluripotent cells, while the abaxial region is farthest from the center of the meristem (Husbands et al., 2009). Leaf primordia emerge in between the adaxial and abaxial regions and develop a clearly defined adaxial side, or upper surface, and an abaxial side, or lower surface. Rapid expansion of the adaxial and abaxial domains result in the boundary formation of leaves, cell type specification, and laminar development. The abaxial-abaxial boundary is the marginal region that separated between the adaxial and abaxial epidermis.

Mature leaves can be further subdivided into regions comprising an epidermis with frequent trichomes and densely-packed lay of palisade mesophyll cells (adaxial), and an epidermis with abundant stomata and loosely-packed spongy mesophyll (abaxial) regions. In the vasculature, the xylem is adaxial to the phloem (Waits & Hudson, 1995). The differentiation of distinctive cell types within adaxial and adaxial domain during leaf development leads to function of gas exchange and regulation of photosynthesis. Adaxialized or abaxialized alterations result in mild to severe defects in leaf formation, trichome distribution, stomatal density and the vasculature organization (McConnell et al., 1998; Eshed et al., 2001; Emery et al., 2003). *HOMEODOMAIN-LEUCINE ZIPPER* (*HD-ZIP*) transcription factors, *KANADI* genes and *microRNA* (*MIR165/166*) are involved in a variety of processes during early development. Within the four classes (I-IV) of *HD-ZIPs, HD-ZIP IIIs* have been shown to have a function in SAM formation, promote adaxial fate, and develop vasculature patterning (Emery et al., 2003; Prigge et al., 2005). In *Arabidopsis*, there are five *HD-ZIP III* transcription factors, *REVOLUTA* (*REV*), *PHABULOSA*(*PHB*), *PHAVOLUTA* (*PHV*), *CORONA*/*ATHB15*, and *ATHB8*. Four of the five *HD-ZIP IIIs* have been shown to regulate adaxial fate in leaf development and are functionally redundant (Prigge et al., 2005). A dominant mutant *phb1-d* altered leaf polarity that developed into a needle-shaped leaf and failed to form a leaf blade (McConnell et al., 1998). On the other hand, *KANADI* (*KAN*) genes promote abaxial fate of leaves (Kerstetter et al., 2001). Loss-of-function of *kan1kan2* double mutants results in adaxialized leaf phenotypes with expanded expressions of *HDZIPIII* (Eshed et al., 2001). Gain-of-function mutants of both genes cause an abaxialized phenotype in the leaf blade and vascular tissue with repressed expressions of *HDZIPIII* (Eshed et al., 2001). *HD-ZIP IIs* have roles in light response, shade avoidance, auxin signaling, and leaf polarity (Ruberti et al., 1991; Ciabelli et al., 2008; Elhiti & Stasolla, 2009; Sessa et al., 2005a, 2018b; Turchi et al., 2013; Merelo et al 2016) . Class II proteins are composed of a homeodomain, an adjacent leucine zipper motif, and a DNA-binding domain (Ruberti et al., 1991). HD-ZIPII proteins contain an LxLxL and CPSCERV motif (Ciabelli et al., 2008; Hermsen et al., 2010). Notably, the LxLxL motif, known as the EAR (Ethylene-responsive binding factor-associated repression) (Ruberti et al., 1991; Ciabelli et al., 2008), has been shown to be important in transcriptional repression in *Arabidopsis* (Kagale and Rozwadowski, 2011). The CPSCERV motif may play a role in sensing environmental cues (Ciabelli et al., 2008). Recently, several HD-ZIP II genes, such as HAT2, HAT3 and ATHb4, were identified that play essential roles in adaxial-abaxial formation (Bou-Torrent, et al., 2012; Turchi et al., 2013; Sessa 2018). The double mutant *hat3 athb4* produced abaxialized and entirely radialized leaves, whereas gain-of-function lines developed up-curled leaves (Bou-Torrent, et al., 2012). Additionally, HAT3 and ATHB4 form a bidirectional repressive circuit to control the balance between adaxial and abaxial fate determination (Merelo et al, 2016). Moreover, some *HD-ZIP IIs*, including HAT1 and ABIG1(also known as HAT22), physically interacted with TPL/TPR in a yeast hybrid assay (Causier et al., 2012). The Groucho/Tup1 corepressor *TPL* and *TPR* families have been implicated in the regulation of diverse developmental processes, including leaf development, hormone signaling, and stress responses (Tao, et al., 2013; Szemenyei et al., 2008; Pauwels et al., 2010). A downstream target analysis of *TOPLESS RELATED 3* (*TPR3*) under drought conditions showed a significant induction by *ABIG1* (Liu et al., 2016).

*KAN* and *REV* have opposite roles in promoting adaxial-abaxial polarity formation (Reinhardt et al., 2013). This antagonistic regulation between *REV* and *KAN* mainly though opposing regulation of downstream targets (Reinhardt et al., 2013). Downstream targets of these transcription factors include class II *HDZIP* (*HD-ZIPII)* transcription factors. Among *HD-ZIP IIs, HAT2* and *ABIG1* were shown to be genes oppositely regulated by *REV* and *KAN* (Reinhardt et al., 2013). Those two genes belong to the Opposite Regulated by *REV* and *KAN* (*ORK*s) genes and are direct targets of *REV* and *KAN1* (Reinhardt et al., 2013; Liu et al., 2016). The *ABA INSENSITIVE GROWTH 1* (*At4g37790, ABIG1*) has been shown to mediate abscisic acid (ABA) growth inhibition, but not stomatal closure. The function of *ABIG1* has been investigated in response to drought stress, but not during plant development (Liu et al., 2016) . Here, we describe a novel function of *ABIG1* in laminar growth and adaxial-abaxial polarity. The *abig1* mutant phenotype exhibits defects in plant growth, leaf formation and patterning. We also provide evidence of the genetic and physical interaction of *ABIG1* with *TOPLESS* (*TPL*), which together regulate leaf development.

## Materials and Methods

### Plant materials and growth conditions

*Arabidopsis* seeds were directly sown in potting medium (ProMix PGX soil mix) or on solid media containing MS Basal Salts (PhytoTech, KS), 0.05% MES, and 0.05% sucrose at, pH 5.7. Seedlings germinated on media were transferred 7-10 days after planting (DAP). Plants were grown in a chamber under 12 h light, 25 °C Day/30 °C night, and ≤50% RH.

### Cloning of constructs and plant transformation

The complete coding sequence of *ABIG1, HAT9*, and *HAT14* was amplified using gene-specific primers (S2 Table). Promoter regions of *ABIG1* were amplified using primers targeting 3-kb upstream sequence. PCR products were cloned into GATEWAY *pENTR/D-TOPO* (Invitrogen, CA) vectors. Entry clones were subcloned into *pMDC32* overexpression (Curtis et al., 2003), *pMDC163 GUS*, and *C/N-YFP* (Bai et al., 2007) destination vectors via LR reaction. Positive clones were verified using PCR and sequencing. *Agrobacterium* (strain: GV3101) transformation was performed by adding 250-500 ng plasmid DNA to 50 μL of thawed electrocompetent cells. The cell suspension was frozen in liquid nitrogen for 5 mins and heat-shock treated at 37 °C for 30 s, returned to ice for 5 mins and shaken at 250 RPM for 3 hours at 28 °C. Bacterial cultures (100-500 µl) were spread onto kanamycin-selective plates and incubated for 2-3 days at 28 °C. Positive colonies were screened using PCR for the corresponding insert size. The constructs *pMDC32:ABIG1* (*35S:ABIG1*), *pMDC32:HAT9* (*35S:HAT9*), *pMDC32:HAT14* (*35S:HAT14*), and *pMDC163:ABIG1* (*pABIG1:GUS*) were introduced into *Arabidopsis* (Col-0) by *Agrobacterium*-mediated transformation as described in (Clough et al., 1998).

### Genotyping and double mutant analysis

For genotyping, genetic crossing and phenotype analysis, the following ecotypes and mutants of *Arabidopsis* (*Arabidopiss thaliana*) were used: Landsberg erecta, Ler, *abig1-1* (ecotype, Ler; Liu et al., 2016), *rev-6* (Ler; Prigge, et al., 2005), *rev-10d* (Ler; (Emery, et all., 2003), *kan1,2,3* (Ler; Eshed et al., 2004), and *tpl-1* (Ler; Long et al., 2002). The genotyping methods and primer sequences for the single and double mutants are listed in S2 Table.

### GUS staining and histology

More than twenty T_3_ transgenic lines were examined for GUS activity. The seedlings, rosette leaves, and inflorescence tissues were harvested and pre-fixed in 80% acetone on ice for 1 hour, followed by submerging into GUS staining solution (100 mM potassium phosphate buffer, pH 7.0, 0.5 mM potassium ferricyanide and potassium ferrocyanide, 10% Tween buffer, and 1 mM X-gluc) at 37 °C for 2 hours or overnight. The chlorophyll was removed by washing stained tissues with 70% ethanol three times. For the vascular tissue observation, all tissues were examined, and photographs were taken using a Nikon stereoscopic dissecting or compound microscope (SMZ1000).

### Leaf sectioning and observation

Ten seedlings and ten rosette leaves were dissected and fixed in 2% (v/v) glutaraldehyde for 2 hours and rinsed in water. All tissues were rinsed, dehydrated through an ethanol series, and embedded in LR white resin (EMS, #14380). Samples were polymerized anaerobically to cure the white resin into transparent capsules. Embedded tissues were trimmed and cut into 2-µm thin sections. The sections were collected and stained with eosin for five min and washed twice, followed by staining with toluidine blue for 30 min before mounting on slides. Slides were examined and photographed using a Nikon compound microscope.

### Scanning electron microscopy (SEM)

Fresh plant tissues were fixed and mounted to the stage of a FEI Inspec-S Scanning Electron Microscope (FEI) and viewed with low pressure and medium scanning speed.

### Quantitative measurement of leaf size and shape

For measurement of the leaf size, shape, and number of cells, the seventh leaf of each WT and mutant plant was cleared in 70% ethanol, flattened, and mounted on slides with cytoseal. The petiole length, blade length, blade width, leaf perimeter, and area were calculated using the macro plugin in Fiji/ImageJ. To measure the cell numbers, three pictures were taken from the tip, middle region near vein, and bottom of the adaxial side of each leaf using a Leica compound microscope and a 10X objective with the same light and contrast settings. The total number of cells was counted in the whole area of every picture. Ten to twelve plants were used for each data point (Maloof et al., 2013).

### Real-Time PCR experiments

Arabidopsis seeds were germinated in liquid medium and grow for 12 days and then treated with dexmethasone for 120 minute or pre-treated with cyclohexmide for 20 minutes. Total RNA was isolated from mock, dexamethasone-treated, cyclohexmide-treated and cycloheximide-dexamethasone treated Col-0, GR-REV and GR-KAN by using Necleospin RNA Plant kit (Macherey-Nagel, www.mn-net.com). cDNA was made from 1 ug total RNA using Tetro cDNA synthesis kit with DNase treatment according to manufacturer’s instructions (Bioline). cDNA was diluted into 100 ul and 2 ul of cDNA was used to perform qRT-PCR. PCR was done using gene-specific primers (see Supplemental Table 1) in technical triplicates on a LightCycler 480 system using the Sensifast SYBR Master mix (Bioline). The ratio of experimental target mRNA to an ACTIN control for each sample was calculated by Applied Biosystems software. An average for the biological replicates and standard deviation were calculated in Excel.

### Yeast two-hybrid assay

The full length *ABIG1, REV* (negative control), *TPL*, and truncated versions of *ABIG1* were amplified from cDNA and sequenced before cloning into GATEWAY® pENTR/D-TOPO® (Invitrogen, CA). The entry clones were subcloned into both pDEST32 and pDEST22 vectors (Invitrogen). The fragments were predicted to produce proteins of various length, designated as ABIG1N (protein sequence, 1-52 amino acids), ABIG1NHD (1-202 aa), ABIG1HD (53-202 aa), ABIG1HDC (53-278 aa), ABIG1C (203-278 aa), and ABIG1 (1-278 aa). To generate the mutated EAR domain in the ABIG1 construct, ABIG1NA (1-52 aa), the <u>L</u>X<u>L</u>X<u>L</u> sequence was replaced with <u>A</u>X<u>A</u>X<u>A</u> by PCR. Yeast two-hybrid interaction assays and color reactions were performed as described in the ProQuest Two-Hybrid System (Invitrogen #PQ10001-01, Carlsbad).

### Biomolecular fluorescence complementation

The coding regions of *ABIG1, REV* and *TPL* were cloned from *Arabidopsis* cDNA as described above. The entry clones were subcloned into pSPYNE*-*35S and pSPYCE*-*35S vectors to generate BiFC constructs for transient expression assays (Walter et al., 2004). The transient tobacco assay methods were modified from (Walter et al., 2004). The tobacco leaves were harvested two days after infiltration and immediately examined using an SP5 confocal microscope (Leica) using the same settings for all samples (gain, contrast and pinhole) to examine the subcellular level of living tobacco cells. This result was consistent in three individual tobacco plants. This experiment was repeated twice.

## Results

### ABIG1 is expressed within the adaxial side of leaves

We have previously demonstrated that *ABIG1* is one of the *ORK* genes (for Oppositely Regulated by *REV* and *KAN*) with a role in ABA-induced senescence (Reinhardt et al., 2013; Liu et al., 2016). Additionally, we also showed that *ABIG1, HAT1*, and HAT2 were regulated by *REV* and *KAN* in dexamethasone (DEX) plus protein synthesis inhibitor cycloheximide (CHX)-treated plants and acted as a direct target for both factors (Liu et al., 2016). Although it was determined that *ABIG1* was up-regulated by *GR-REV* (Glucocorticoid Receptor, GR) and down-regulated by *GR-KAN1*, little is known about the role of *ABIG1* during the establishment of tissue adaxial-abaxial polarity. Initially, we performed Real-Time PCR expression analysis to confirm all the HDZIP II family members in regulation of *REV* and *KAN* and found that the majority of them are regulated by *REV* and *KAN* (Fig. 1A, and Fig S1). Thus, to elucidate the polarity function of *ABIG1*, we investigated its expression in all plant tissues throughout plant development (Fig. 1B-G). We generated a promoter construct that includes the *ABIG1* coding region and 3-kb upstream, fused to the reporter β-glucuronidase (GUS) gene, and transformed it into *Arabidopsis Col-0*. At the seedling stage (7 days), GUS staining showed strong *ABIG1* expression in the vascular tissues of cotyledons, leaf primordia, roots and inflorescences (Fig. 1B-E). To further determine whether *ABIG1* expression was also associated with leaf polarity development, London Resin (LR)-white sections were made of seven-day-old GUS reporter seedlings and leaf primordia from 10 individual T2 lines. Longitudinal and cross sections of the shoot apex revealed that GUS expression was strongest in the leaf primordia, lateral stipules, and the adaxial side of epidermal cells in young and mature leaves (Fig. 1B, C, F). In the rosette leaves, the expression was found in both adaxial and abaxial epidermal and mesophyll cells but stronger in adaxial epidermis (Fig. 1G). These observations showed that *ABIG1* was active in the adaxial sides of leaf in the seedling stage and expanded to mesophyll and abaxial side of leaves in the mature stage, suggesting a role for *ABIG1* in leaf development and adaxial-abaxial polarity patterning during leaf expansion. During reproductive development, we observed that *ABIG1* expression was also present in young floral buds, petals, and filaments (Fig. 1E), suggesting a potential role during reproduction. This result is in agreement with a previous finding that *ABIG1* was involved in floral organ polarity development (Shchennikova et al., 2018).

**Fig 1.**
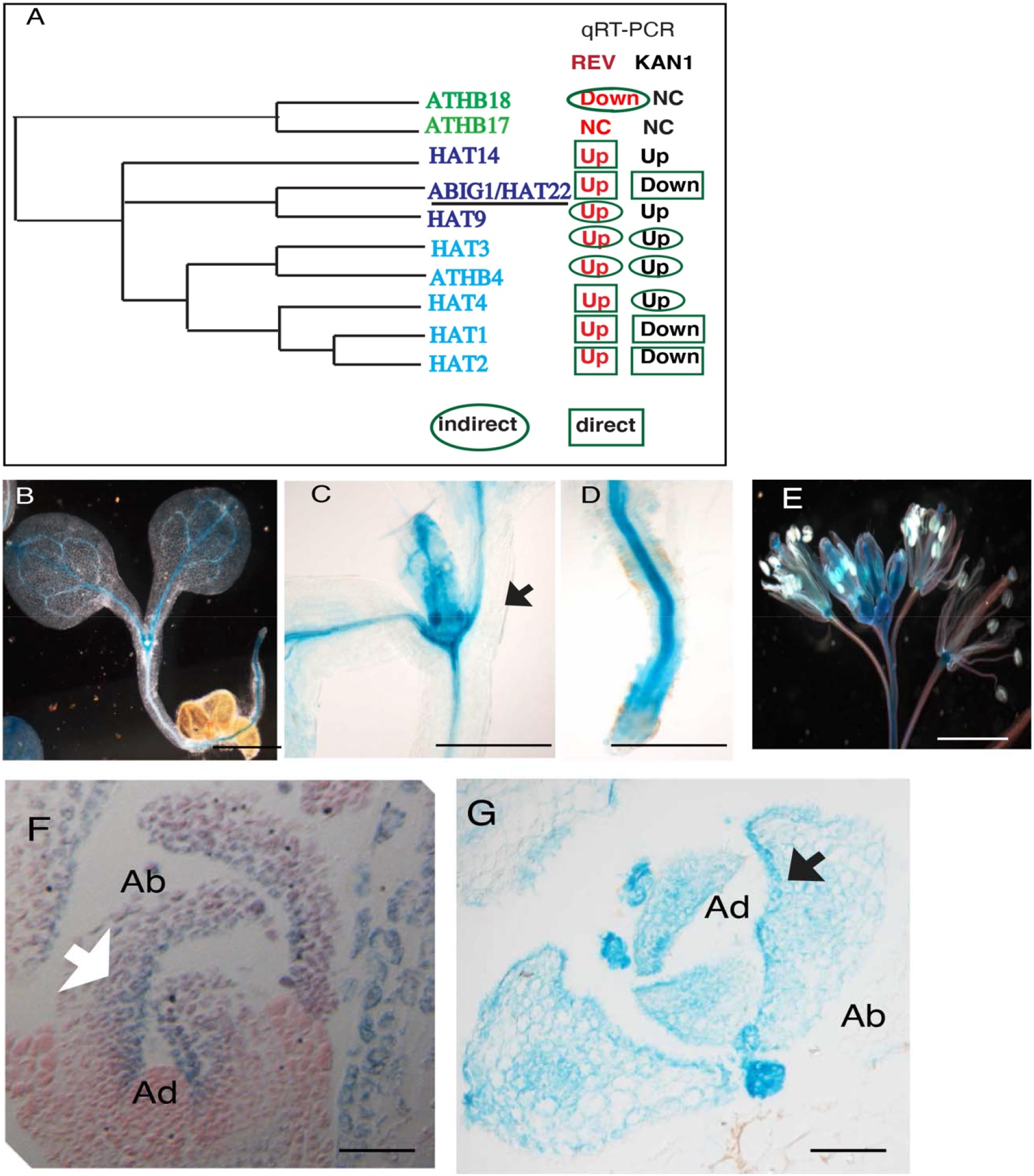
Expression analysis of *ABIG1*. (A) Schematic representation of the HDZIP II genes regulated by *REV* and *KAN*. Green boxes indicate direct target, green cycles indicate indirect targets. ‘Up’ indicates up-regulation, ‘Down’ indicates down-regulation. NC, indicates no significant changes. (B) Gus staining expression analysis in *pABIG1::GUS*-transformed *Arabidopsis* plants. Five-day-old seedlings showing expression in the vascular tissue, Bars = 2 mm. (C) cotyledons, (D) root, and (E) inflorescence. Bars = 2mm for B, Bar=50 μm for C, D, and E. (e) Longitudinal section of the shoot apex at 7 days showing leaf primordia. (F) Cross-section of young and (G) mature leaves in the shoot apex. Bars = 100 μm for F-G.

### Ectopic expression of *ABIG1* created adaxial-abaxial polarity defects

In order to gain further insights into the function of *ABIG1*, we generated *ABIG1*-overexpression lines using the CaMV 35S promoter (*35S:ABIG1*) in the Col-0 background. The overexpression of *ABIG1* resulted in severe growth defects in both the T1 and T2 generations (Fig. 2). Compared to WT, 12-day old of *35S:ABIG1* seedlings were smaller with extremely narrower leaf blades and up-curled leaves (Fig. 2B). Scanning electron microscopy (SEM) images showed that the narrow leaf blade and up-curling phenotypes of *35S:ABIG1* occurred early during the emergence of the true leaves (Fig. 2D). Later in plant development, when new leaves were fully expanded, the leaves of the overexpression lines remained curled upward towards the adaxial leaf surface, in contrast to the flattening of wild-type leaves (Fig. 2E-F). When transplanted to soil, *35S:ABIG1* plants remained dwarfed and displayed small narrow leaf blades and up-curled leaves (Fig. 2G, M, O). In addition, there was no internode elongation, and a very short stem developed in the *35S:ABIG1* plants. Upon flowering, *35S:ABIG1* plants typically produced abnormal inflorescence with few flowers that were sterile (Fig. 2G). In addition to the phenotypic observation, we performed quantitative measurement of the leaf size and shape to determine the differences between WT and the *35S:ABIG1* line (Fig. 2H-K). In the *35S:ABIG1* plants, there was a significant decrease in petiole length, blade length and width (Fig. 2H) as well as in blade perimeter and area (Fig. 2I-J). Furthermore, examination of the cell size in true leaves from 12-day-old plants (12 plants, the seventh leaf per plant) revealed that *35S:ABIG1* had a greater number of cells in the tip, middle, and bottom regions of the leaves (Fig. 2K). This analysis revealed that overexpression of *ABIG1* resulted in phenotypes that affect polarity along with reduced lamina development, thereby decreasing leaf size. This is consistent with previous observations showing that establishment of adaxial and abaxial patterns is required for leaf blade formation (Bowman et al., 2002; Lin et al., 2003).

**Fig 2.**
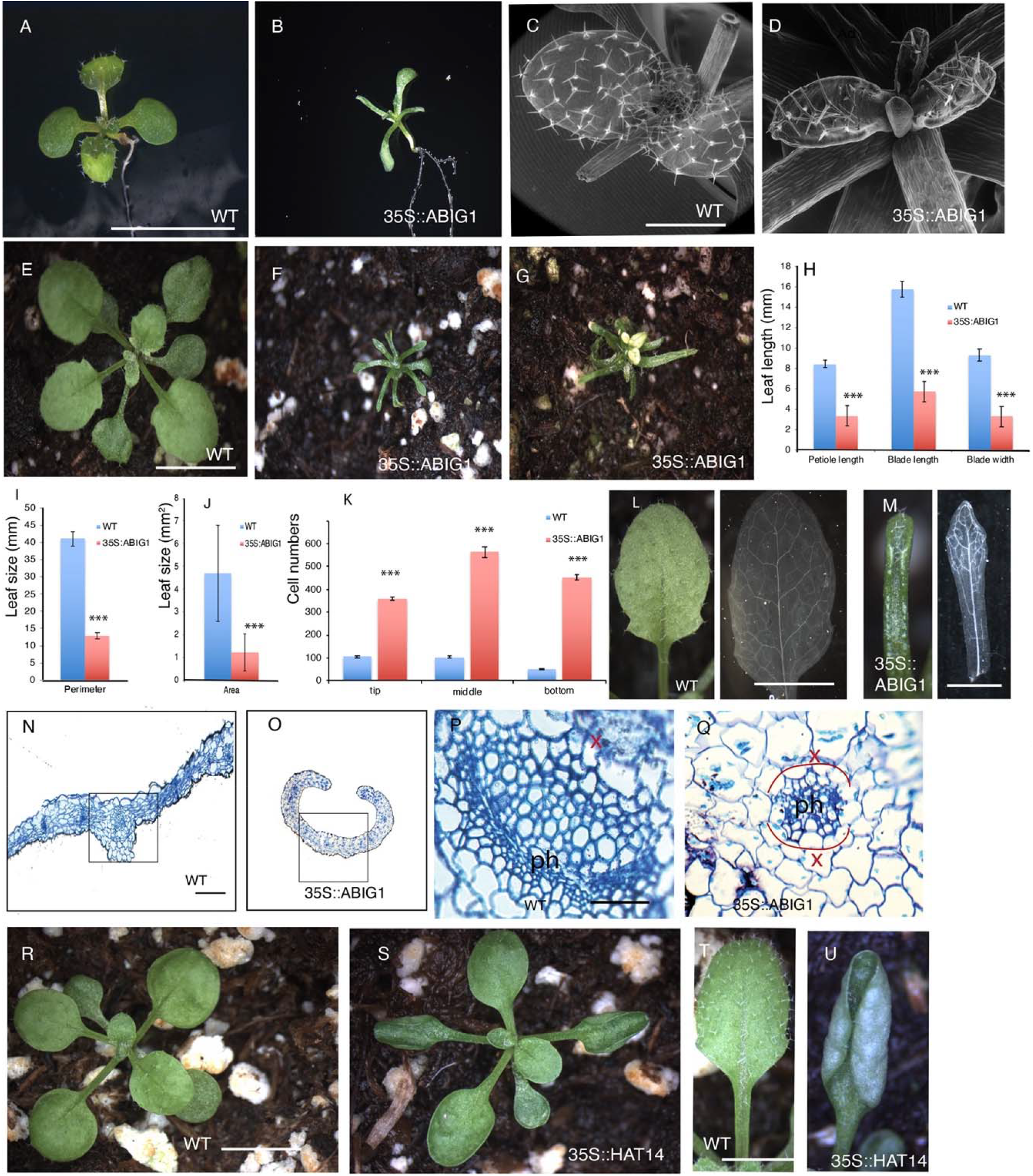
Overexpression of *ABIG1* induces Adaxial-abaxial Polarity Defects. Phenotype of (A) WT and (B) dwarf *35S::ABIG1* seedlings. Scanning electron microscopy of (C) WT and (D) *35S::ABIG1* narrowed leaf blade and leaf primordia showing up-curled phenotype, Bars = 50 μm. (E) WT and severe dwarf 35S:ABIG1 of (F) mature plants and (G) reduced inflorescence. (H-K) Quantitative measure of leaf size and shape in WT and *35S:ABIG1* 12-day-old seedlings. The y-axis shows the length of true leaf length (H), leaf perimeter (I), area (J) and cell numbers (k). Measurements were calculated relative to the wild type in three independent biological experiments, ^***^*P<0*.*001* by Student’s *t*-test. Wild type plants with normal vein pattern in cleared rosette leaves (L), flat leaf blade (M), and vascular bundle organization (P). (P) and (Q) are close-up images for (N) and (O) in the black box area, respectively. *35S::ABIG1* showed narrow leaf blade and abnormal vein pattern in cleared leaves (M), extreme up-curled leaf blade (O), and disorganized vascular bundle (Q). Bars = 50 μm for M and N. Bars = 10 μm for O and P. Phenotype of (S) showed up curled leaves in *35S::HAT14* seedlings and (U) rosette leaves. Bars = 50 μm for M and N. Bars = 10 μm for O and P.

Examination of cleared, fully expanded rosette leaves revealed that the venation pattern was not interrupted (Fig. 2M). To further examine the phenotypic defects in the vasculature of *35S:ABIG1* plants, mature leaves were collected, sectioned, and stained with toluidine blue. In the vascular bundle, transverse sections through the midvein of *35S:ABIG1* leaves displayed an altered vascular pattern (Fig. 2N-Q). In contrast to the distinct adaxial-abaxial polarity patterning of the vasculature in WT plants (Fig. 2N, P), the organization of the phloem and xylem cells were disrupted in the *35S:ABIG1* plants (Fig. 2O, Q). The phloem tissues were surrounded by xylem cells, a phenotype similar to those of the *kan1kan2kan3* triple mutant or a dominant gain-of-function *rev-10d* mutant (Emory, et al., 2003).

To further investigate the *ABIG1* involvement in polarity formation and leaf development, we overexpressed two other closely related HDZIPIIs, *HAT9* and *HAT14*. We were not able to detect any obvious growth defect in *35S:HAT9*, however, we observed a phenotype on the leaf development of the *35S:HAT14* mutant (Fig. 2S, U). The phenotype in gain-of-function of *HAT14* was similar to but less severe than that of *ABIG1*. Sixteen out of 20 transgenic lines showed up-curled leaves in seedling stage (Fig. 2S). Consistent with up-curled leaf phenotype in *35S:ABIG1*, those plants developed unusual phenotype with up-curled leaf blade in rosette leaves (Fig. 2U). These results showed a clear relationship between *ABIG1* and *HAT14* in leaf polarity development.

### Polarity-mediated leaf development requires *ABIG1* functions

The mutant line *abig1-1*, an enhancer trap line in the *Landsberg erecta* background and a knock down allele of *abig1* mutant, has been reported to be insensitive to ABA treatment and tolerant to drought stress (Liu et al., 2016). The enhancer trap system uses Ac/Ds transposable elements and a GUS reporter gene to identify the expression pattern and regulatory cascade of the trapped gene (Springer, 2000). To characterize the roles of *ABIG1* in adaxial-abaxial pattern formation, the *abig1-1* mutant was examined during leaf development (Fig. 3). At the seedling stage, the true leaves of homozygous *abig1-1* plants displayed a subtly downward curled phenotype (20/20, Fig. 3) compared to WT (0/20, Fig. 3A). The observed phenotype was consistent with the previously described *revoluta* phenotype (Reinhart et al., 2013). Since mRNA levels of *ABIG1* are up-regulated by *REV* and down-regulated *KAN*, we decided to obtain higher order mutants. *REV* is a positive regulator of *ABIG1* expression and has been shown to induce a number of *HD-ZIP II* genes including *HAT3* (Reinhart et al., 2013). The *rev-6* mutant displays a slightly downward curled leaf phenotype (22/24, Fig. 3C). In the homozygous *rev-6 abig1-1* double mutant, the leaf down-curling phenotype of the rosette leaves was more severe than the *rev-6* and *abig1-1* single mutants (15/20, Fig. 3D). In contrast to the relatively weak *rev-6* allele, we found around 25% of the double mutant significantly affected shoot development (Fig. 4B-F). The phyllotaxy of the floral branches in *rev-6 abig1-1* mutant was irregular, and the length of the internode was often reduced (Fig. 4B). Additionally, the double mutant exhibited a stronger phenotype in reproductive growth and development, as the *rev-6 abig1-1* produced a radialized, bladeless cauline leaf, suggesting the loss of adaxial identity in emerging leaf primordia (Fig. 4E-F). In *rev-6 abig1-1*, a single flower formed at the flower primordia and developed into a single silique (24/30, Fig. 4C) indicating that the floral primordia were arrested at an early developmental stage. Additionally, several entirely radialized axillary buds (Fig. 4E-F) were observed in *abig1-1 rev-6*. The phenotype was consistently observed to the next generation with high frequency (12/12). We did not observe this phenotype with radialized axillary buds in the *rev-6* mutant (22/22). These data indicated that loss of *ABIG1* function enhanced the downward curled phenotype of the *rev* mutant that resulted in extremely abaxialized phenotype.

**Fig 3.**
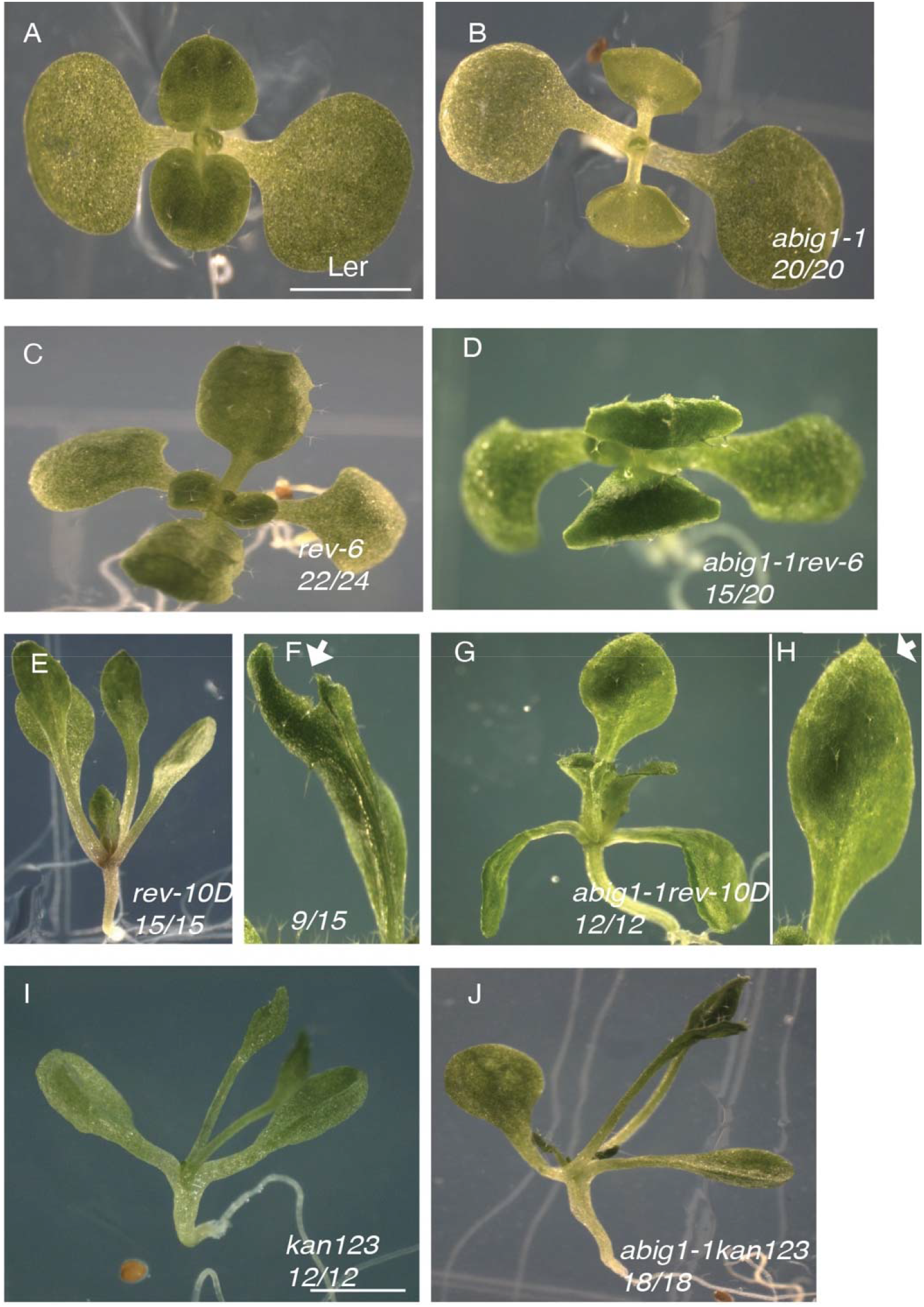
Mutations in ABIG1 and REV affect leaf adaxial formation. (A) Wild-type and (B) enhancer trap *abig1-1* mutant that showed down-curled leaves. (C-D) Leaves of a *rev-6* and a *rev-6 abig1-1* double mutant exhibited downward leaf curling, with the double mutant showing enhanced phenotypes. (E-H) A *rev-10d, rev-10d abig1-1* double mutant exhibited upward leaf curling phenotypes. (I-J) Leaves of *kan123* triple, *kan123abig1-1* quadruple mutant. Bars = 5mm.

**Fig 4.**
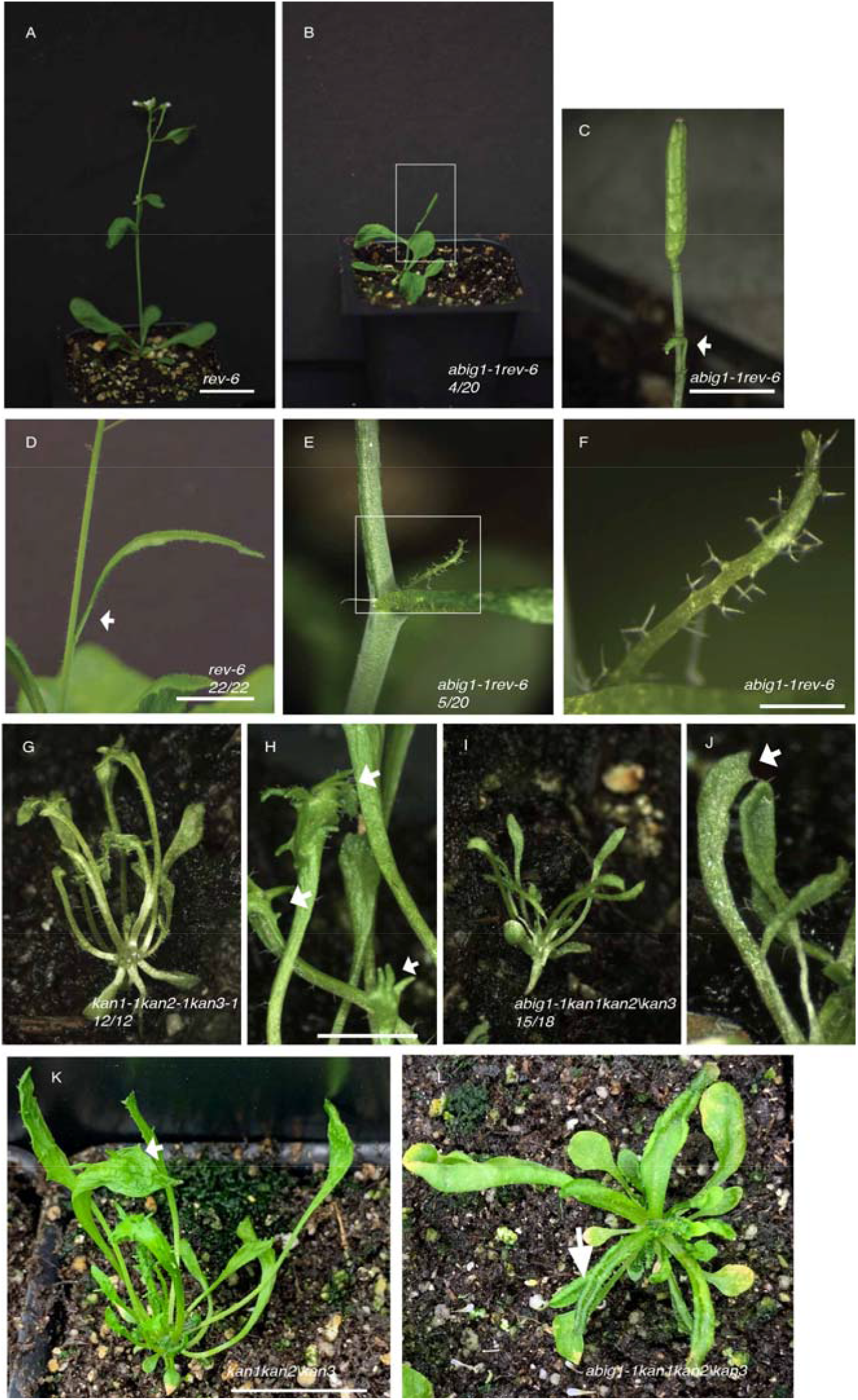
The genetic interaction between ABIG1 and KAN-REV signaling. (A) *rev-6* mutant. (B-C) *abig1-1 rev-6* showed outgrowth of arrested inflorescence with terminated single silique. White arrowhead indicated the arrested axillary bud. (D-F) double mutant displayed radialized leaf-like structure in cauline leaves. Bar = 1cm for A-E, Bar = 0.5cm for F. (G) *kan1-2kan2-1kan3-1* mutant, (H) a close-up image of leaf finger in *kan1-2kan2-1kan3-1* mutant. (I) *abig1-1kan1-2kan2-1kan3-1* mutant, (J) a close-up image of *abig1-1kan1-2kan2-1kan3-1* displayed smooth leaf blade. Bar = 1cm in (G)-(J). Bar = cm in (L)-(K).

We hypothesized that *REV* and *ABIG1* function together in establishing adaxial patterning. To address this possibility, we generated a double mutant between *abig1-1* and a gain-of-function *REV* mutant, *rev-10d* (Emery et al., 2003). In *rev-10d* mutants, upward curled (15/15, 15 plants out of 15 plants) and fused leaves (9/15, 9 plants out of 15 plants) are often observed (Fig. 3E-F). The *rev-10d abig1-1* double mutant caused reduced phenotype in contrast to *rev-10d* single mutant (Fig. 3G-H). The seedlings of *abig1-1 rev-10d* produced rosette leaves that are flat and less curled upward (12/12) (Fig. 3H), the frequency of fused leaves was also significantly reduced (0/12, Fig. 3G-H). Overall, these observations indicated that loss-of-function of *abig1-1* mutant reduced *rev-10d* phenotype indicating *ABIG1* primarily contributes to adaxial identity.

*KAN* is a negative regulator of *ABIG1* expression (Reinhart et al., 2013). To more clearly define the function of *ABIG1* in adaxial polarity establishment, we crossed *abig1-1* mutants to the polarity defective triple *kan1kan2/+kan3* knockout mutant, in which the adaxial-abaxial polarity in most leaves is severely disrupted that forms long petiole and upward curled leaves (Fig. 3I). The quadruple mutant (*abig1/kan1/kan2/kan3*) did not exhibit additional defects in leaf polarity in the seedling stages, displaying long petioles and narrow leaf blades (Fig. 3J). Once transplanted to soil, the *kan1kan2kan3* triple mutant remained dwarf but also developed a severe phenotype with lobed leaves and a leaflet-like structure growing out of the leaf blade (12/12, Fig. 4G-H). In contrast to the triple mutant, the *abig1-1kan1kan2kan3* quadruple mutant had few leaflet-like structures on the leaf blade and a partial reduction of the extreme adaxialized phenotype in the *kan1kan2kan3* triple mutant (15/18, Fig. 4I-J). This phenotypic difference was more obvious at maturity where the *abig1-1kan1kan2kan3* quadruple mutant displayed upcurled leaves without leaflet-like tissue on leaf blade (18/18, Fig. 4K). This suggested that loss of *ABIG1* function caused by the abig1-1 mutation partially rescue the severe adaxialization phenotype exhibited by the *kan1kan2kan3* triple mutant.

### Genetic interaction between *ABIG1* and *TOPLESS*

To further analyze how *ABIG1*is involved in leaf development and regulates leaf polarity patterning, we examined the genetics and biochemical interaction between ABIG1 and its downstream target. We previously performed RNA-sequencing of an estradiol-induced *ABIG1* line (*XVE:ABIG1*) and identified a small number of downstream targets (Liu et al., 2016). The majority of targets are involved in stress-responsive pathways such as ABA and JA signaling pathways (Liu et al., 2016). The data indicated that *TOPLESS RELATED 3* (*TPR3*) was significantly induced by *ABIG1* (Liu et al., 2016). Since the role of TPL/TPR corepressors were involved in various developmental processes. We focused on investigating genetic and biochemical interaction between ABIG1 and TPL/TPR. As the TPL/TPR loss-of-function mutants show no obvious phenotype due to functional redundancy, we explored the genetic interaction between *abig1-1* and *tpl-1*, a dominant-negative mutant that is involved in embryonic development (Smith & Long, 2010) by generating double mutants of *abig1-1* and *tpl-1. tpl-1*, a temperature-sensitive mutant (Long et al., 2002). When grown at 22 °C, *tpl-1* mutants displayed pin-shaped, fused, and cup-shaped cotyledons (Fig. 5C-E). The *abig1-1 tpl-1* phenotype appeared to be enhanced compared to *tpl-1*, in that 3% of the double homozygous seedlings (6/188) failed to form a shoot and grew two roots when germinated at 22 °C (Table S1; Fig. 5F-G). The shoot-to-root transformation phenotype was only observed in the *tpl-1* single mutant when embryos developed at 29 °C (Long et al., 2002). The number of single cotyledons in the *abig1-1 tpl-1* mutant (34%, 63/188) was higher than in the *tpl-1* single mutant (20%, 38/192) (Table S1). Later in development, seedlings that began with single and fused cotyledons continued developing cup-shaped true leaves (Fig. 5I), a phenotype never observed in the *tpl-1* mutant (Fig. 5H). At flowering, double mutants formed dwarf plants, smaller than *abig1-1* and *tpl-1* single mutants (Fig. 5L). These results suggested that the *abig1-1 tpl-1* double mutant enhanced the *tpl* mutant phenotype. Furthermore, *ABIG1* expression in *abig1-1 tpl-1* seedlings was distinguishable from *abig1-1* in that GUS staining was found in the vasculature of double-root seedlings, cup-shaped cotyledon seedlings, fused-cotyledons, and at the topmost region of pin shaped seedlings in the double mutant (Fig. 5G, J-K). These genetic results indicated that *ABIG1* is necessary for leaf formation in *tpl-1* mutant plants.

**Fig 5.**
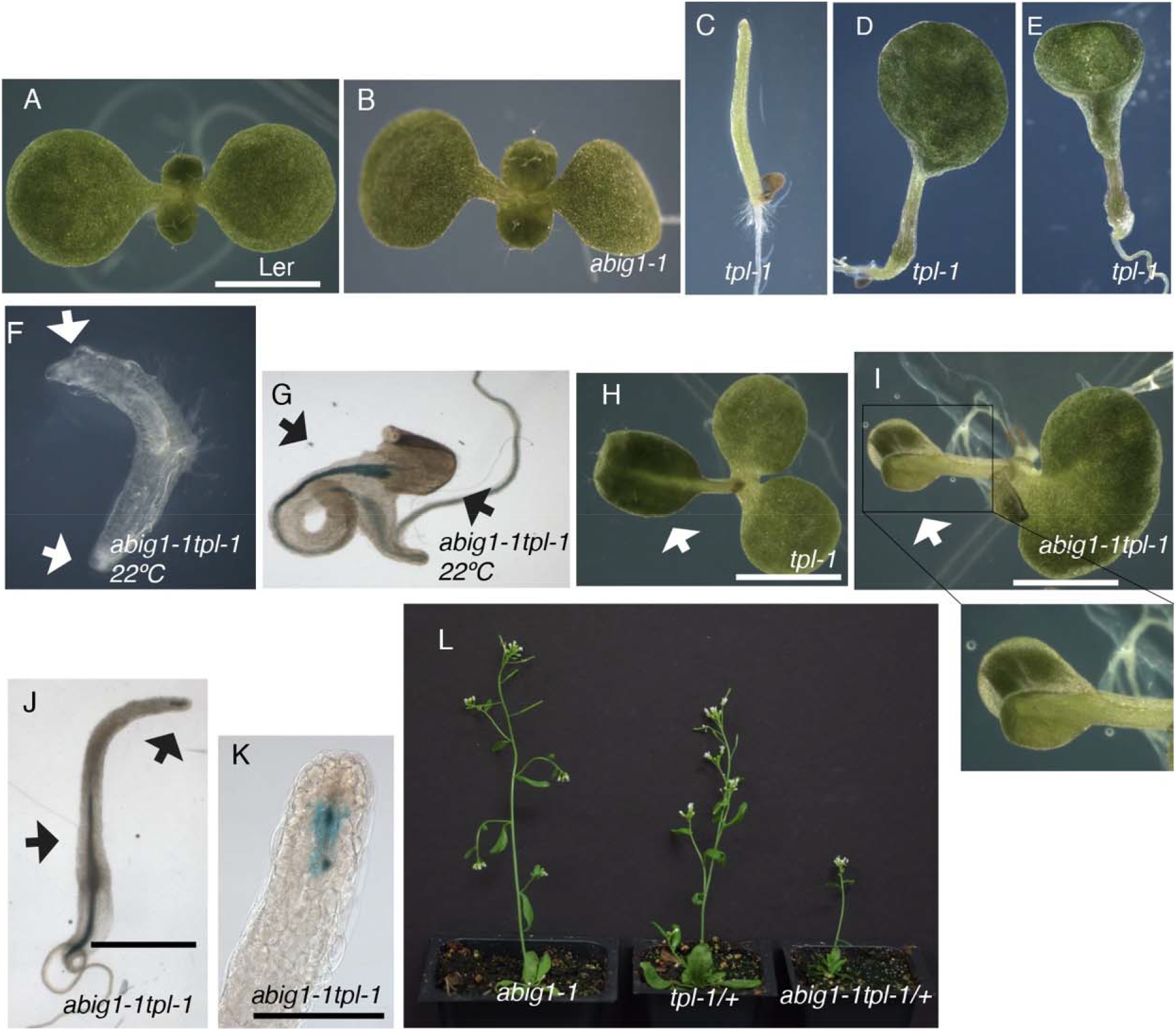
*TPL* genetically interacts with *ABIG1*. (A) WT, and (B) *abig1-1* seedlings at 10 days after germination. (C-E) The *tpl-1* mutant displayed pin-shaped, single, and fused cotyledon phenotype at the permissive 22 °C. (F) A true leaf emerged from a fused cotyledon in the *tpl-1* mutant. White arrowhead indicated flatten leaf in tpl-1 mutant. (F-G) The *abig1-1 tpl-1* double mutant had an enhanced phenotype. White arrowheads indicate shoot-to-root phenotype in the double mutant. Double roots were observed when grew at 22 °C. (I) A cup-shaped true leaf formed in the *abig1-1 tpl-1* mutant. (J-K) GUS expression pattern of *abig1-1* in *abig1-1 tpl-1* mutant. Black arrowheads indicate GUS expression. (L) *abig1-1 tpl/+* was a dwarf mutant compared with the single mutant. Bar = 5mm.

### ABIG1 physically interacts with TPL corepressors

*TPL* has been shown to act as a corepressor (Causier et al., 2012). Corepressor recruitment is largely governed by the presence of an EAR (Ethylene-responsive binding factor-associated repression) motif. To test the hypothesis that ABIG1 and TPL physically interacted through the EAR domain, we initially generated a variety of truncated ABIG1 variants to characterize the key regions of ABIG1 required for interaction with TPL (Fig. 6A) to test protein-protein interactions using a yeast two-hybrid assay (Fig. 6A). The GAL4 binding domain (BD) was fused to TPL as bait, and the GAL4 activation domain (AD) was fused to ABIG1 as prey. An interaction between the two proteins occurred in yeast (Fig. 6B). Yeast two-hybrid analysis of a full-length, unaltered REVOLUTA*-MEK (Magnani & Barton., 2011), Fig. 6A, construct 8), an HD-ZIP III protein lacking an EAR motif, showed no β-gal activity. Next, the ABIG1 domains were analyzed for their necessity for interaction with TPL using seven truncated ABIG1 configurations (Fig. 6B, constructs 1-7), including a mutated EAR motif (LXLXL to AXAXA, 2). Constructs 1, 3 and 7 contained the minimal N-terminus region required for transcriptional activation of β-gal, and each harbored the EAR motif. Construct 2, carrying a mutated EAR motif, did not interact with ABIG1. This indicated that the EAR domain of ABIG1 could strongly interact with TPL.

**Fig 6.**
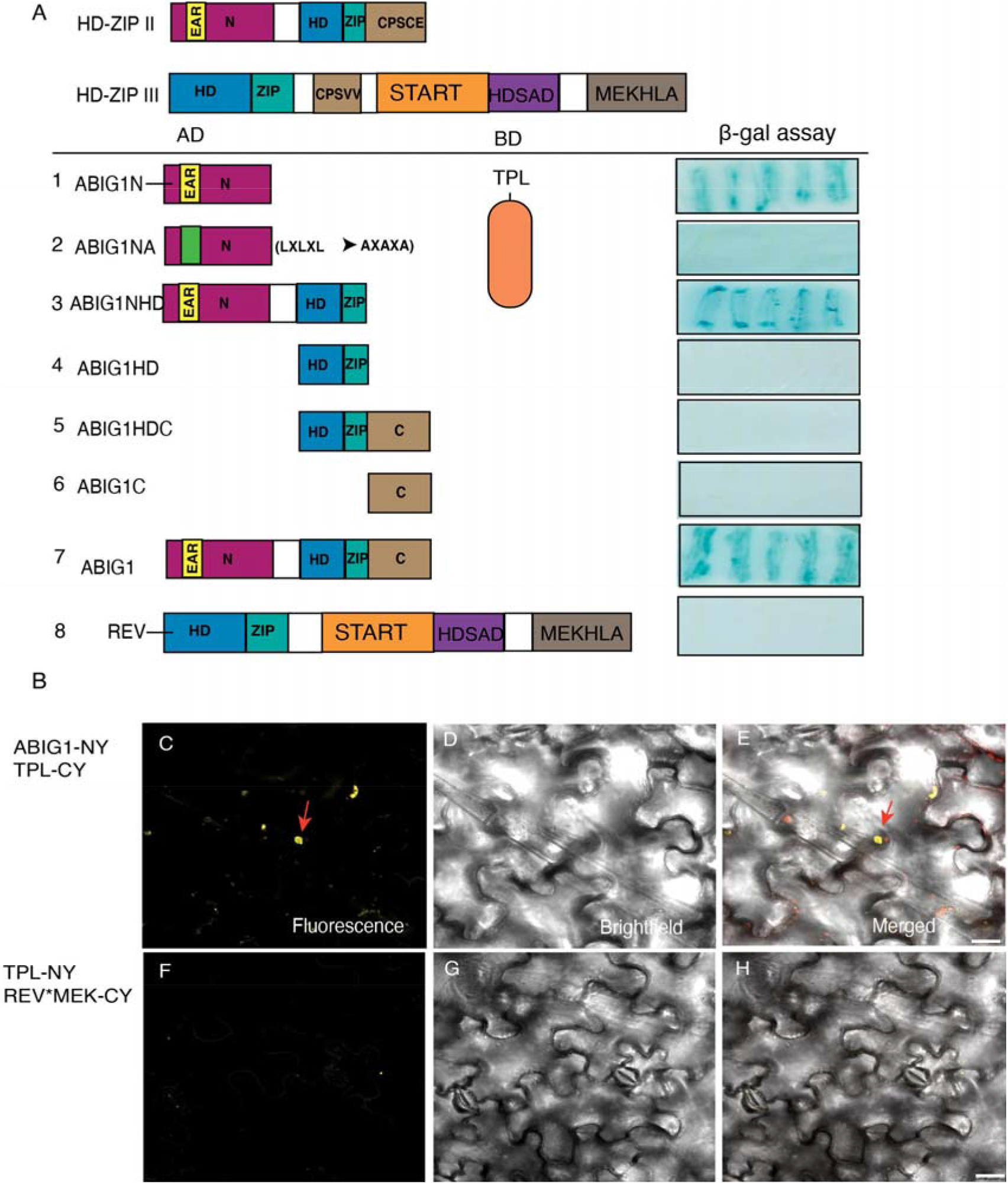
ABIG1 interacts with TPL *in vitro* and in vivo. (A) Yeast-two-hybrid assay to test the interaction between full-length or truncated versions of ABIG1 and TPL proteins. *B*-gal assays were used to indicate the interaction. Constructs 1-7 present the different truncated versions of the ABIG1 protein; 1, ABIG1N (amino acid, 1-52); 2, ABIG1NA (aa, 1-52; <u>L</u>X<u>L</u>X<u>L</u> changed to <u>A</u>X<u>A</u>X<u>A); 3,</u>ABIGNHD(aa, 1-202); 4, ABIGHD (53-202); 5, ABIG1HDC (53-287); 6, ABIG1C (203-287); 7, full-length ABIG1; and 8, REV as a negative control. (B) BiFC assays were used to test the ABIG1 interaction with TPL in *N. benthamiana* leaves. Yellow dots and red arrowheads indicate a positive signal. There was no signal detected from the negative control REV^*^-MEK. Bars = 10 μm.

To confirm whether ABIG1 interacts with TPL *in vivo*, we carried out bimolecular fluorescence complementation (BiFC) analysis (Fig. 6B). ABIG1 was fused to the N-terminus of YFP, and TPL was fused to the C-terminal end. Consistent with yeast two-hybrid results (Fig. 6A), there was an interaction between ABIG1 and TPL, and the signal was clearly evident in the nucleus (Fig. 6B). A negative control using REV*-MEK C-terminus paired with N-terminus TPL lacked a fluorescent signal. Together, these two assays suggested that ABIG1 interacts with TPL both *in vitro* and *in vivo*.

## Discussion

We previously demonstrated that *ABSCISIC ACID INSENSITIVE GROWTH 1 (ABIG1)* is oppositely regulated by *REV* and *KAN* (*ORK*) genes and that *ABIG1* can inhibit leaf growth and leaf production at the shoot apical meristem and can retard root growth and cause leaf yellowing under drought conditions (Liu et al., 2016). The aim of this paper was to investigate the role of *ABIG1* in establishing lateral organ polarity in *Arabidopsis*. Our study presents evidence to support the hypothesis that *ABIG1* is necessary for laminar growth and leaf polarity establishment in *Arabidopsis*. Firstly, plants carrying a mutation in *ABIG1* show defects in leaf polarity formation that caused a down-curled phenotype in rosette leaves. When *ABIG1* was overexpressed, plants exhibited upwardly curled leaves, defects in leaf size and shape, and altered vascular patterning. Secondly, *pABIG1:GUS* expression was restricted to the adaxial side of leaf primordia and the expanded mature leaf. Thirdly, the genetic interaction within a double *ABIG1* and *REV* mutant showed a severely abaxialized phenotype with downwardly curled leaves and an enhanced radial-shape to cauline leaves. Other double mutants, namely *abig1-1rev-10D* and *abig1-1kan1,2,3*, rescued the leaf curling and polarity phenotype. These results suggested that *ABIG1* functions in laminar growth and similarly to *REV* as an adaxial regulator and plays a role in adaxial polarity formation. Therefore, we suspected that *ABIG1* may be involved in other regulatory networks that control leaf laminar growth and adaxial-abaxial fate decision. Among the potential targets of *ABIG1*, we investigated *TPL* using genetic and biochemical approaches to test any interactions. Similar genetic interactions between *TPL* and *REV* have been reported (Smith and Long., 2010). We found that *abig1 tpl-1* double mutants showed enhanced double-root phenotype. The majority of HD-ZIP II proteins, including ABIG1, have an EAR domain in their N-terminal region, indicating they have may act as transcriptional repressors. We examined the interaction of ABIG1 with the corepressor TPL using Yeast Two-Hybrid and Bi-molecular Fluorescence Complementation (BiFC) assays. Our results showed that ABIG1 and TPL directly interacted through the EAR domain in both yeast and tobacco epidermal cells. Our data show that *ABIG1* plays a role in leaf laminar development in *Arabidopsis*.

### *HD-ZIP II* genes are required for adaxial-abaxial polarity establishment in *Arabidopsis*

Among the ten members of the *HD-ZIP II* gene family in *Arabidopsis, ATHB2, HAT3* and *ATHB4* function redundantly in establishing the dorsal-ventral axis in cotyledons and developing leaves (Bou-Torrent, et al., 2012). . It is worth noting that *HAT3* and *ATHB4* are expressed in the abaxial domain of leaf and genetically interact with the HD-ZIP III genes *REV, PHB* and *PHV* to control both SAM development and bilateral symmetrical patterning during leaf formation (Tuchi et al., 2013). In addition, *JAIBA (JAB)/HAT1*, a close homolog of *HAT3* and *HAT2*, regulates meristematic activity during formation of different tissues and organs during both vegetative and reproductive stages (Zuniga-Mayo et al., 2012). In agreement with these findings, we found that the *HD-ZIP II ABIG1* is involved in adaxial fate determination, as has been found for other *HD-ZIP II* genes such as *HAT1, HAT3, ATHB2*, and *ATHB4* (Tuchi et al., 2013; Merelo et al., 2016). . Although *ABIG1* is not within the same subfamily as *HAT1, HAT3* and *ATHB4*, it is in a sister clade (Ciabelli et al., 2008). There are two additional *HD-ZIP II* genes within this subfamily, *HAT9* and *HAT14*, which are the closest relatives to *ABIG1* in *Arabidopsis* (Ciabelli et al., 2008). Given the ability of *HD-ZIP II* genes to negatively auto-regulate and the close homology, *HAT9* and *HAT14* may also contribute to *ABIG1* functions. A similar leaf curled upward phenotypic change was observed in *35S:HAT14*, suggesting that *HAT14* also involved in leaf polarity formation. Additionally, the observation that both *HD-ZIP II* and *III* genes are involved in abaxial-abaxial polarity development might indication that HD-ZIPs function in the same pathways. Based on the genetic and biochemical interactions between HD-ZIP II and III genes and proteins, functional redundancy was observed during apical formation and leaf polarity establishment in embryos (Tuchi et al., 2013). However, it is still unclear whether *HD-ZIP II* genes act independently in organ polarity or if their activities depend upon the *HD-ZIP IIIs*.

### ORK genes play dual roles in hormone-mediated development and stress responses

*ABIG1* is regulated by *REV* and *KAN. REV* and *KAN* display opposite roles in establishing leaf adaxial-abaxial polarity (Reinhart et al., 2013; Liu et al., 2016). *REV* acts as an activator and *KAN* acts as a repressor in the transcriptional response to leaf boundary formation. They antagonize each other by regulating downstream targets in opposite directions, such as a couple of transcription factors that play regulatory roles in various hormone-mediated developmental and stress-responsive processes. Brandt et al., (2012) has previously shown that the expression of another *ORK* gene, *TAA1*, an auxin biosynthesis enzyme, is activated in the adaxial domain by *REV* while repressed in the abaxial side by *KAN*. Another pair of *ORK* genes, *PYL6* and *CIPK12*, are related to ABA responses. *PYL6* is an ABA receptor (*RCAR/PYR1/PYL*) that binds to ABA and leads to inactivation of the *SNF1-Related Kinases* (*SnRKs*) that regulate the ABA signaling pathway (Qin et al., 2008). CIPK12 is a member of the SnRK3 family that is involved in ABA-dependent signaling and polarized pollen tube growth (Steinhorst et al., 2015). Little is known about how ABA signaling is involved in leaf polarity. However, *PYR1*, which is required for ABA transport, displays differential adaxial-abaxial expression patterns, which is directed by *mir165/166* and their target HD-ZIP III (Yang et al., 2019). Similarly, the *ABIG1* gene plays a role in mediating endogenous ABA signals and in response to drought treatment. We noted that *ABIG1* is also increased by drought in floral tissues (Su et al., 2013). This is particularly interesting because we hypothesize that the genes oppositely regulated by *REV*/*KAN* may have a dual role in both development and stress responses. Indeed, Song et al., (2016) discovered that *ABIG1* appeared to be a ‘hub’ gene for the ABA-responsive pathway that may regulate gene expression in all aspects of ABA-related processes. However, it has yet to be determined if ABA-regulated formation of adaxial-abaxial patterning occurs through *mir165/166* and an *HD-ZIP III*. Much less is known about how ABA is involved in regulation of *KAN* and its transcriptional repression during seed germination and leaf development. As such, more detailed studies will be needed to determine the interactions among *HD-ZIP IIIs, HD-ZIP IIs*, and *KAN* in mediating ABA action during early meristem development, polarity formation, and stress responses. Therefore, it will be interesting to reveal if additional regulatory networks integrate responses by which leaf development can be adjusted in response to the changing environment.

### The evolutionary relationship among *HD-ZIP II* genes hints at ancient roles in leaf development and stress responses

The HD-ZIPI, II, III, and IV proteins are remarkably convergent within the HD and ZIP domains, with other domains distinguishing each HD-ZIP gene family. HD-ZIP I and II genes are usually involved in environmental signaling responses. HD-ZIP II members contain a unique seven amino acid motif (CP-X-CER-X) at the carboxy terminus, which may be responsible for the interaction with other proteins that sense environmental signals. Genome-wide analysis in *Medicago truncatula* showed that the HD-ZIP genes had distinctive, tissue-specific patterns and divergent responses to various stresses (Li et al., 2020). In rice, *small grain and dwarf 2 (SGD2)*, encoding an *HD-ZIP II* transcriptional repressor, controls plant leaf and panicle development by regulating gibberellin biosynthesis (Chen et al., 2019). Thus, it is quite possible that the tissue-specific regulation of *HD-ZIP II* genes could balance growth and stress responses.

Six out of ten HD-ZIP II proteins in *Arabidopsis* include an EAR-motif (LXLXL) at the amino terminus that is crucial for transcriptional repression. Consistent with this hypothesis, the HD-ZIP II protein HAT1 forms a transcriptional repression complex with TPL to inhibit target genes that modulate the anthocyanin biosynthesis pathway (Zheng et al., 2019). The molecular mechanisms by which those HD-ZIP IIs bind to TPL to repress downstream transcriptional activities in development and stress responses remain to be identified. In *Eucalyptus camaldulensis*, overexpression of a homologue of *ABIG1, EcHB1*, results in increased density of cells and chloroplasts in leaves, a higher photosynthesis rate, as well as drought tolerant features (Sasaki et al., 2019). It will be interesting to see if EcHB1 can physically interact with EcTPL.

Understanding how *HD-ZIP II* genes function in leaf development will give us a more complete view of the role of these genes throughout the plant kingdom. As the expression levels of these genes are influenced by environmental factors (red light or drought, depending on the family member) and these genes interact with a major developmental pathway, the *HD-ZIP II* genes are excellent candidates as hubs for integrating intrinsic developmental pathways with environmental signaling pathways.

## Acknowledgements

Financial support from the National Science Foundation in America is gratefully acknowledged. A small part of the data was collected in the lab of Dr. Kathryn Barton at the Carnegie Institute for Science. We thank Kathy Barton and the Barton lab members for providing technical support and comments for the manuscript.

## Data availability statement

All data generated or analyzed during this study are included in this published article and its supplementary information files.

## Supplementary data

**Fig S1. Class II HD-ZIPs are in response to 60-min dexamethasone treatment with and without cycloheximide as measured by qRT-PCR**. The relative expression values are means of three biological replicates and three technical replicates. Arabidopsis seeds were germinated in liquid medium and grow for 12 days and then treated with dexmethasone for 60 minute or pre-treated with cyclohexmide for 20 minutes. PCR was done using gene-specific primers (see Supplemental Table 2) in technical triplicates on a LightCycler 480 system. The ratio of experimental target mRNA to an ACTIN control for each sample was calculated by Applied Biosystems software. An average for the biological replicates and standard deviation were calculated in Excel. ‘++’ indicates dexamethasone and cyclohexmide treatment. ‘--’ indicates mock control without any treatment. A. HDZIPIIs in response to 60-min dexamethasone treatment with and without cycloheximide as measure were regulated by REV, B. HDZIPIIs were regulated by KAN.

**Table S1. *ABIG1* enhances the *tpl-1* mutant phenotype**. Value are means of three replicates SD. Statistical significance of the new phenotype in the *Student t-test* (*P* <0.05) is represented by an asterisk.

**Table S2. Primer sequences for genotyping and cloning**.

